# Microfluidic Droplet-Assisted Single-Cell Raman Sorting for High-Throughput Screening of Functional Lignin-Degrading Microbes

**DOI:** 10.1101/2025.04.25.650636

**Authors:** Tingting Jia, Manling Zhang, Huicheng Chen, Ling Lu, Lin Wang, Yunhua Wang, Guoxia Zheng

**Author notes:** Corresponding Author E-mail addresses. These authors contribute equally (Tingting Jia, Manling Zhang).

## Abstract

Lignin, a major component of plant biomass, poses significant challenges for sustainable valorization due to its structural recalcitrance. Traditional microbial screening methods are hindered by culturability limitations and time-consuming workflows. This study introduces a novel Microfluidic Droplet-assisted Single-Cell Raman sorting system (MD-SCR sorting) to address these bottlenecks. By integrating droplet microfluidics with single-cell Raman spectroscopy (SCRS), the platform enables non-destructive, high-throughput screening of lignin-degrading microbes directly from environmental samples. A deuterium tracing strategy was developed, where microbial strains were cultured in lignin media containing 30% D_2_O. Raman detection of C-D bond vibrations (2040–2300 cm^−1^) allowed in situ identification of active degraders within 12 hours. The MD-SCR system achieved precise single-cell encapsulation, cultivation, and sorting using a custom-designed three-layered microfluidic chip. Validation experiments demonstrated strong correlation (R^2^=0.92) between %C-D values (quantifying deuterium assimilation) and traditional shake-flask degradation efficiency. Two lignin-degrading fungi (Pleurotus ostreatus G5 and P. eryngii PB) exhibited distinct C-D peaks, while non-degrading controls showed no deuterium incorporation. Laser intensity optimization (5% power) ensured spectral fidelity with >95% cell viability. The platform’s label-free, culture-independent workflow facilitates rapid functional screening of unculturable microbes, overcoming limitations of fluorescence-based methods. This technology not only advances lignin bioconversion by enabling targeted isolation of high-efficiency degraders but also establishes a scalable framework for exploring complex microbiomes. By bridging metabolic phenotyping with single-cell resolution sorting, the MD-SCR system offers transformative potential for sustainable resource utilization and environmental biotechnology.

## 1 Introduction

Plant biomass, the most abundant renewable carbon resource on Earth, is regarded as an ideal feedstock for bioenergy and bio-based chemicals. Composed primarily of lignin, cellulose, and hemicellulose, lignin alone constitutes 10-30% of lignocellulosic biomass, making it the second most abundant natural organic polymer in nature [1, 2]. This complex, amorphous, three-dimensional phenolic network exhibits remarkable structural stability and serves as a critical component of cell walls in woody plants. However, its inherent heterogeneity and recalcitrance to degradation have led to its classification as an industrial byproduct [3]. The composition of lignin varies significantly across sources (softwood, hardwood, or herbaceous crops) [4], with annual global production estimated at 5×108 to 36×108 tons *5+. In bioethanol production, lignin is generated as a major byproduct [6], while agricultural advancements have also yielded vast amounts of lignin-rich straw residues, representing a largely untapped resource [7]. Furthermore, the pulp and paper industry produces “black liquor,” an alkaline waste containing lignin, phenolics, and other pollutants. Despite constituting only 10-15% of total wastewater, black liquor accounts for 90-95% of environmental toxicity due to its lignin-derived compounds [8, 9]. Globally, 40-50 million tons of lignin are annually generated by this industry, yet merely 1.5% is commercially utilized, primarily as low-value lignosulfonates [10].

The recalcitrant structure of lignin poses significant challenges for its valorization. Efficient decomposition and conversion of lignin are critical for sustainable resource utilization, environmental protection, and renewable chemical production. While lignin conversion research has advanced since the 1980s [11], current depolymerization methods—including thermochemical oxidation, hydrogenolysis, gasification, and hydrolysis—remain energy-intensive and costly [12]. In contrast, microbial biocatalysis offers a promising alternative due to its specificity and operational efficiency under mild conditions [13, 14]. However, traditional microbial screening methods rely on culture-dependent techniques, which are time-consuming (months-long cycles) and fail to access >99% of unculturable soil microorganisms [15, 16]. Developing culture-independent strategies for in situ functional microbe identification and high-purity isolation is thus imperative for sustainable lignin management.

Single-cell screening technologies have revolutionized microbial studies by bypassing cultivation requirements. Over the past decade, advanced tools such as optical/magnetic tweezers, dielectrophoresis, flow cytometry, microfluidics, and single-cell Raman spectroscopy (SCRS) have emerged as pivotal platforms [17]. SCRS, a label-free and non-invasive technique, captures biochemical fingerprints of individual cells by detecting photon energy shifts from molecular vibrations [18]. Its unique capability to track metabolic activity via stable isotope probes (e.g., ^13^C, D_2_O) enables in situ identification of functional microbes. Isotopic incorporation into biomolecules induces characteristic Raman band shifts, allowing metabolic quantification at single-cell resolution [19]. Nevertheless, environmental complexity, laser-induced cell damage, and technical limitations in achieving precise single-cell sorting hinder SCRS applications [20, 21].

To address these challenges, droplet-based microfluidics has emerged as a powerful tool for high-throughput single-cell manipulation. By encapsulating individual cells into picoliter-scale droplets, this technology creates isolated microreactors that preserve native metabolic states while enabling precise analysis and sorting. However, existing platforms lack automated, non-destructive workflows for in situ cultivation and functional screening. This study integrates microfluidic droplet technology with SCRS to develop a Microfluidic Droplet-assisted Single-Cell Raman sorting system (MD-SCR sorting). A droplet-based microfluidic chip was engineered to achieve high-throughput single-cell encapsulation, forming bio-microreactors for in situ cultivation. Combined with Raman detection, this system enables accurate identification of lignin-degrading microbes in complex environmental samples. A deuterium tracing strategy was established by culturing lignin-degrading strains and two non-degrading controls in lignin media containing 30% D_2_O. Raman detection of C-D bond signatures (2040-2300 cm^−1^) after 12 h incubation enabled non-destructive identification of active degraders. The %C-D values correlated strongly with traditional shake-flask degradation data (R^2^=0.92), confirming method reliability. This integrated platform achieves high-throughput, non-destructive screening of functional microbes, advancing lignin valorization and offering a novel paradigm for exploring environmental microbiomes. The findings not only deepen understanding of lignin bioconversion but also provide a scalable technological framework for sustainable resource utilization.

## 2 Materials and Methods

### 2.1 Microfluidic Chip Design and Operation

We developed a three-layered microfluidic chip integrating single-cell capture, in situ culture, and Raman detection (Fig.1). The top layer features a 50 μm-diameter microwell array (30 μm depth, 1 mm spacing) for single-cell isolation, optimized via fluid dynamics simulations to maximize capture efficiency and minimize fluid disturbance. A chromium-coated middle layer enhances Raman signal-to-noise ratio by suppressing background interference, while the transparent glass substrate facilitates optical imaging. The chip was fabricated by soft lithography as previously described.

**Figure 1.**
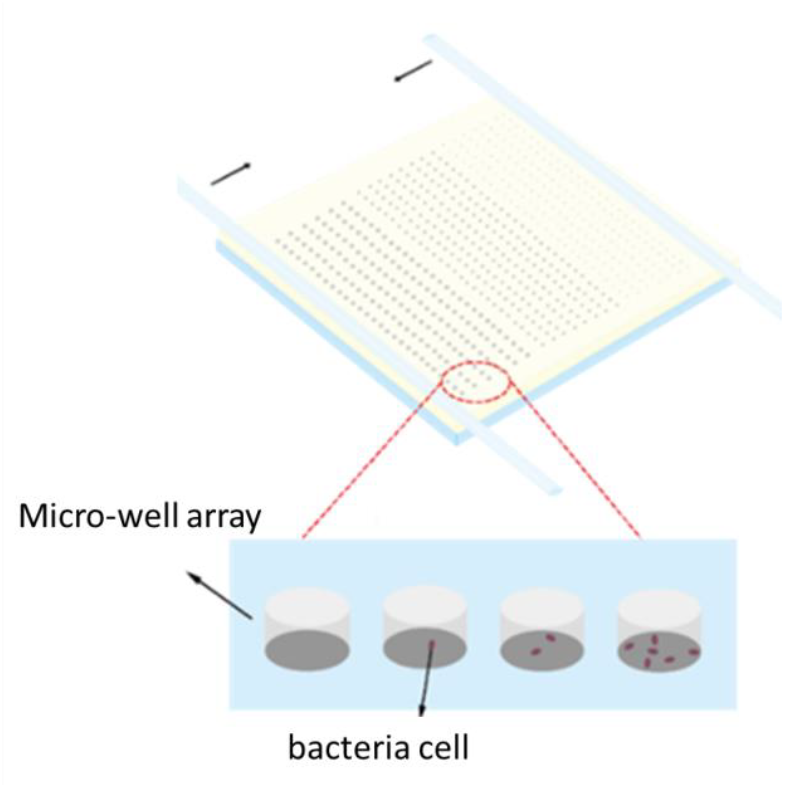
Schematic diagram of microfluidic structure

For operation, deuterium-labeled samples were centrifuged, diluted (∼200 cells/mL), and loaded onto the chip. Single-cell encapsulation was achieved via Poisson distribution-guided scraping. Raman detection utilized a 532 nm laser (5% power, 600 grating) with 5-s acquisitions across 800-3200 cm^−1^. Post-calibration with silicon standard (520.7 cm^−1^), 20 cells/sample were analyzed. Data processing (LabSpec6/Origin2021) included baseline correction and C-D peak quantification (2040-2300 cm^−1^). High-activity cells were selectively cultured in microwells with fresh medium for expansion.

### 2.2 Microbial Cultivation

Lignin-degrading strains (*Pleurotus ostreatus* G5, *P*.*eryngii* PB) and two non-degrading controls (*Enterobacter sp*. NC02, *Kluyvera sp*. pb29) were obtained from Dalian University. Fungi were cultured on PDA/bacterial strains on LB agar. Liquid cultures utilized nutrient broth (bacteria) or Sabouraud medium. Degradation experiments employed nitrogen-limited media (fungi) or modified mineral salt media (bacteria). All media were autoclaved (121°C, 20 min) before use.

### 2.3 Raman Spectroscopy Optimization

Laser intensity effects were evaluated using *P. eryngii* PB. Raman spectra (532 nm laser, 600 grating) were acquired at 0.01-25% laser power. Signal-to-noise ratios and peak resolution analysis identified 5% intensity as optimal, balancing spectral quality with minimal photodamage.

### 2.4 Lignin Degradation Assay

Inoculated nitrogen-limited media (250 rpm, 30°C) were sampled every 24 h. Alkali lignin degradation was quantified via UV-Vis absorbance at 280 nm:

Degradation (%) = (A_0_ − At)/A_0_ × 100%

where A_0_ and A_t_ represent initial and timepoint absorbance.

### 2.5 Deuterium Incorporation Analysis

Cultures were supplemented with D_2_O (0-50% v/v) in 24-well plates. Growth (OD600 for bacteria; ultrasonicated fungal biomass) was monitored every 4 h. For Raman analysis, cells were harvested (8,000 rpm, 10 min), washed twice with deionized water, and air-dried. Single-cell spectra (800-3200 cm^−1^) were acquired using 5% laser power (100× objective). C-D incorporation was quantified as %C-D = [C-D/(C-D + C-H)] × 100% via LabSpec6/Origin2021.

## 3 Results

### 3.1 Optimization of Raman Laser Intensity

Laser intensity optimization is critical for balancing spectral quality and cellular integrity in microbial Raman spectroscopy. Using *Pleurotus eryngii* PB as a model, we systematically evaluated seven laser intensities (0.01%, 0.1%, 1%, 3.2%, 5%, 10% and 25%) with a 532 nm excitation source, 100× objective, and 800-3200 cm^−1^ spectral range. Samples were air-dried and calibrated using silicon’s 520.7 cm^−1^ peak. Each intensity condition was tested with 10 replicate scans per sample. Data processing included baseline correction and normalization using LabSpec6 and Origin2021.

At 0.01% intensity, cells maintained structural integrity but produced no detectable Raman signals. Increasing to 0.1% yielded faint C-H peaks (2800-3000 cm^−1^) with persistent background noise. Optimal signal-to-noise ratios emerged at 1-5% intensity, where the morphological characteristics of microbial cells remained unaltered. Within this intensity range, the peak intensities in Raman spectra increased proportionally with laser power. At 10% laser intensity, optical microscopy revealed no cell lysis or thermal ablation, yet Raman spectra exhibited marked spectral anomalies, indicating laser-induced alterations to intracellular molecular architecture. When intensity reached 25%, cells displayed distinct photothermal damage characteristics (membrane disruption, carotenoid degradation) accompanied by pronounced spectral modifications.

Integrating morphological observations with spectral data, the 5% laser intensity enabled acquisition of well-resolved characteristic peaks while preserving >95% cell viability. This optimized condition effectively mitigated molecular structural damage associated with high-intensity irradiation while enhancing spectral signal quality through excitation energy optimization, thereby establishing robust experimental parameters for quantitative metabolic phenotyping in subsequent microbial analyses.

### 3.2 Strain Isolation and Heterogeneity Analysis

Raman spectral analysis revealed significant fingerprint peak variations among lignin-degrading strains, indicative of distinct metabolic profiles and/or cellular structural divergence. Comparative morphological studies demonstrated conserved colony morphologies across strains under both native environmental conditions and glucose-defined media. However, Raman spectra exhibited marked differences in intracellular metabolite profiles, particularly in lignin-related vibrational bands.

The morphological stability of strains under carbon-limited conditions—regulated by core genetic programs—suggests evolutionary adaptations to lignin-rich habitats. Notably, Raman fingerprint variations provide a non-destructive analytical tool for assessing strain diversity, with spectral signatures serving as auxiliary indicators for functional classification. This morphological conservation across laboratory and natural environments highlights the robustness of cellular architecture despite metabolic plasticity. These findings elucidate molecular diversity and environmental adaptability within ligninolytic microbial consortia, establishing a framework for targeted screening of high-efficiency degraders. Meanwhile, many questions, such as biochemical identification of differential spectral peaks and their correlation with degradation efficiency and dynamic effects of environmental stressors on strain morphology-spectral phenotype coupling, deserve further investigation.

In Raman spectroscopic detection, the 600-1800 cm^−1^ range serves as the characteristic fingerprint region for microorganisms, effectively reflecting the composition and structural features of intracellular macromolecules (Fig.3). This spectral region encompasses vibrational information from critical biomolecules such as proteins, nucleic acids, and lipids. Microbial specificity arises from variations in the proportional composition and spatial configurations of these components, forming unique spectral fingerprints. The complexity of microbial fingerprint regions manifests not only through characteristic signals of individual components but also through synergistic vibrational interactions among multiple biomolecules. This multidimensional spectral information combination provides a robust chemometric foundation for microbial classification, facilitating the development of rapid identification systems based on Raman spectral features. Preliminary assignments of detected characteristic peaks reveal that fingerprint variations among the strains primarily originate from differences in biochemical composition: Fungal signatures are distinct bands at 750 cm^−1^, 1128 cm^−1^, and 1583 cm^−1^, corresponding to cytochrome C vibrational modes, while bacterial signatures are prominent peaks at 1240 cm^−1^ (C-O-P-O-C in RNA backbone) and 1480 cm^−1^ (Amide II bands) (Fig.3).

**Figure 2.**
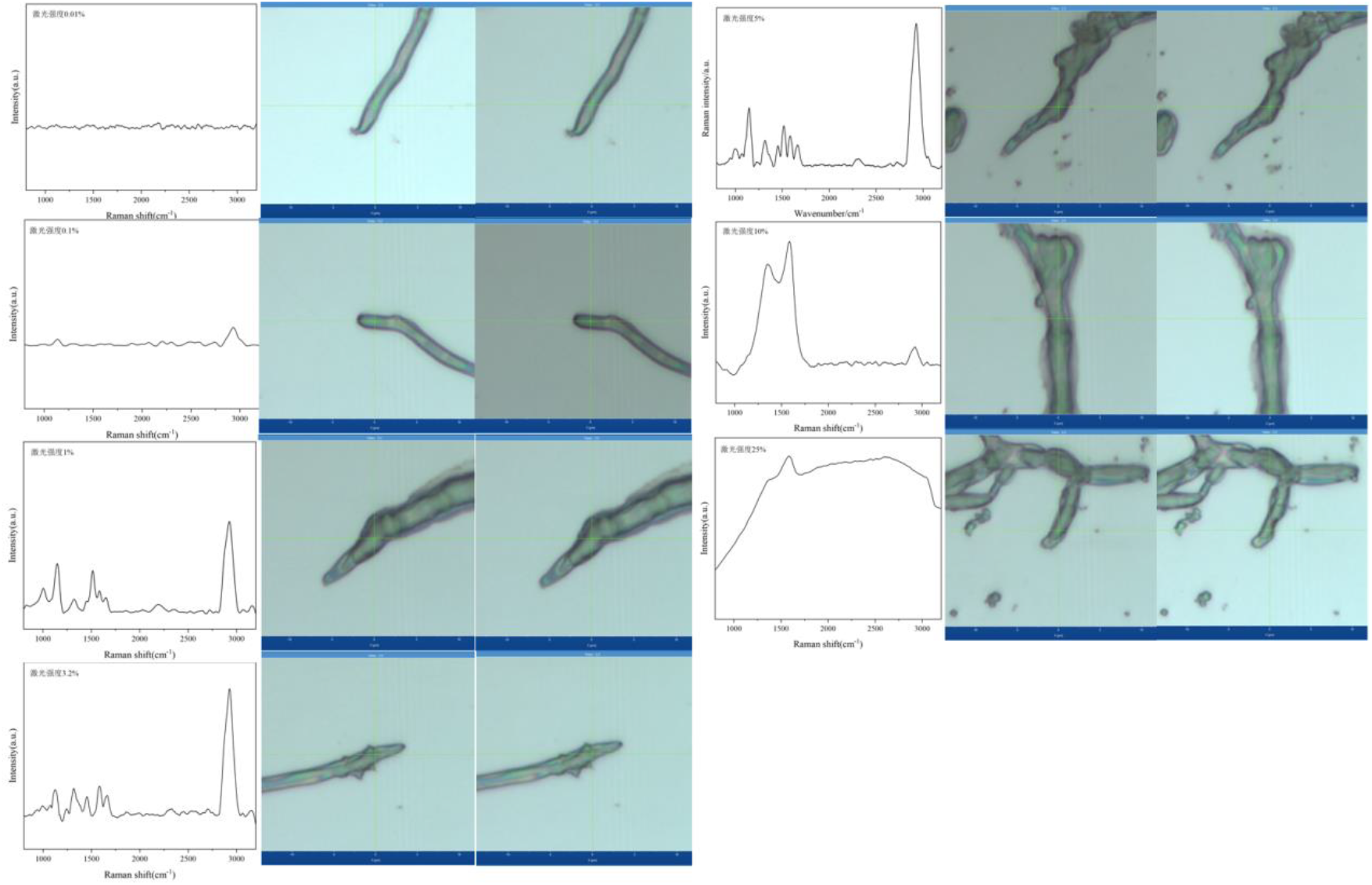
Raman spectra and morphology of PB at seven laser intensities

**Figure 3.**
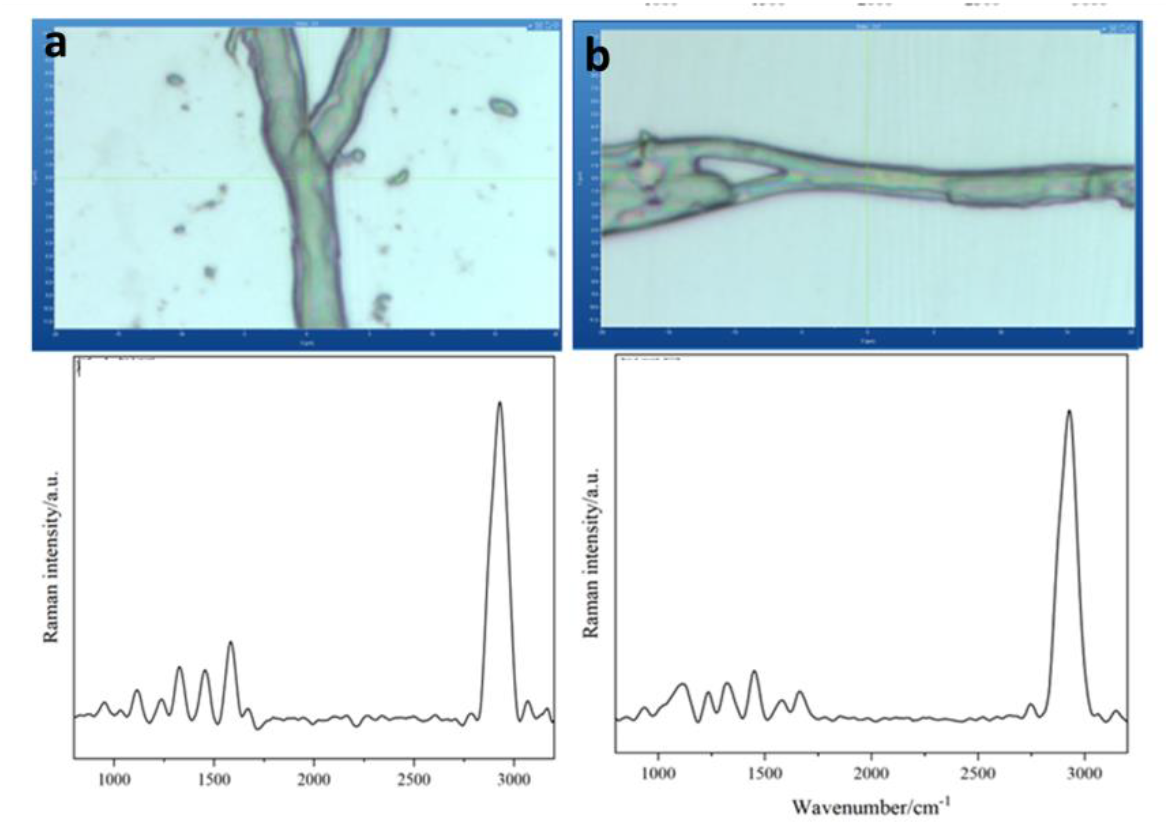
Morphology and Raman spectra of (a) *Pleurotus ostreatus* G5 and (b)*P*.*eryngii* PB in the in situ environmen

Raman spectral characteristic peak analysis enables efficient strain identification [22]: All fungal strains exhibit characteristic peaks at 1308 cm^−1^ (L-phenylalanine) and 1455 cm^−1^ (uracil). The 1660 cm^−1^ peak (L-valine), common across strains, serves as a fundamental diagnostic marker. Additionally, the 1004 cm^−1^ peak (L-phenylalanine) is prevalent in PB and other strains. Combinatorial analysis of these characteristic peaks provides precise criteria for rapid strain discrimination, demonstrating critical application value in distinguishing closely related strains or confirming specific species.

### 3.3 Growth and Deuterium Assimilation of Lignin-Degrading Strains under Different D_2_O Concentrations

Previous studies have demonstrated varying effects of D_2_O on microbial growth, including inhibition, stimulation, or no impact depending on concentration [23–25+. To select a D_2_O concentration without growth inhibition for subsequent experiments, lignin-degrading strains were cultured in liquid expansion media containing 0%, 10%, 30%, or 50% D_2_O. Optical density changes were monitored over 24 h.

Compared to the 0% D_2_O control, low concentrations (10%, 30%) showed no significant growth inhibitionto both strains. In contrast, 50% D_2_O exhibited strain-dependent inhibitory effects: *P. eryngii* PB displayed mild inhibition between 12–20 h (Fig.4a), while *P. ostreatus* G5 showed pronounced suppression after 8 h (Fig.4b). Since 30% D_2_O caused showed slight growth inhibition in both lignin-degrading strain, this concentration was selected for further experiments.

**Figure 4.**
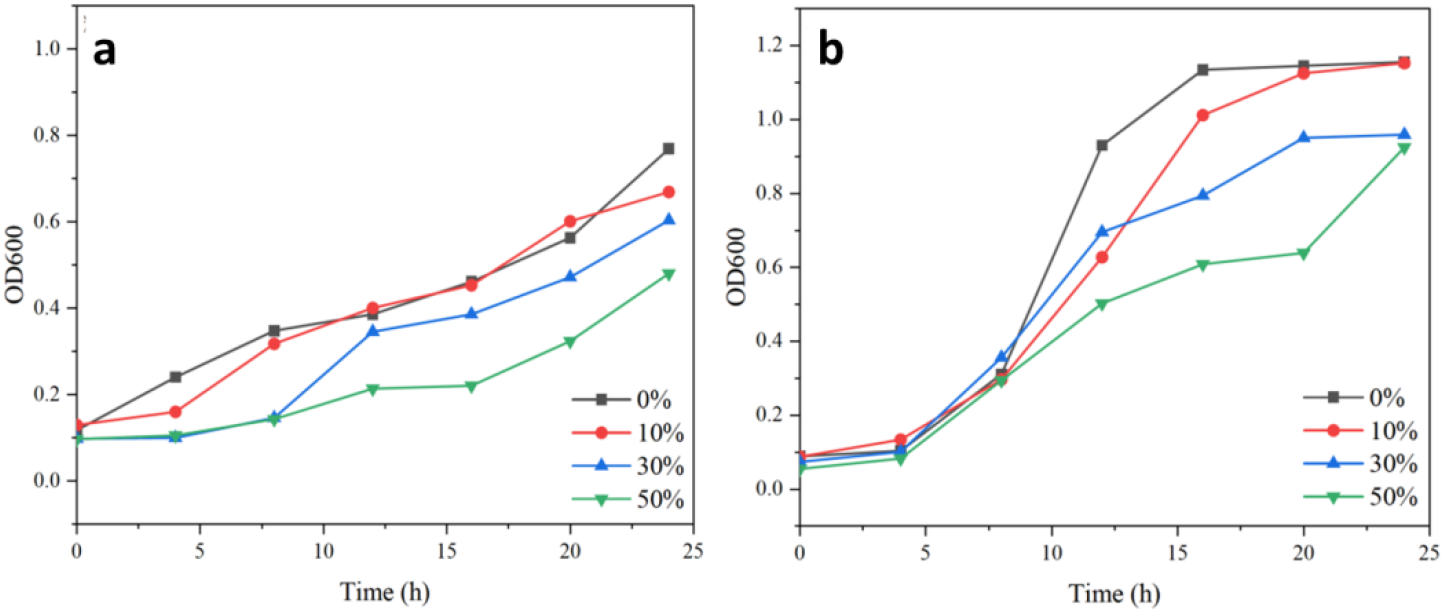
Growth curves of (a) *Pleurotus ostreatus* G5 and (b)*P*.*eryngii* PB with different D_2_O concentrations

### 3.4 Investigating Lignin Utilization Activity of Lignin-Degrading Strains via D_2_O Labeling

Four test strains—lignin-degrading G5 and PB, and non-degrading controls pb29 and NC02—were cultured in lignin degradation media containing 30% D_2_O with alkali lignin as the sole carbon source. Raman spectroscopy (800–3200 cm^−1^ range) was employed to detect intracellular deuterated carbon (C-D bonds), focusing on the 2040–2300 cm^−1^ region.

As shown in Fig.5, lignin-degrading strains exhibited distinct C-D peaks (centered near 2170 cm^−1^) in D_2_O media(Fig.5a,b), while controls showed only baseline noise in this region(Fig.5c,d). Non-degrading strains (pb29, NC02) displayed no C-D signals in D_2_O media, with spectral profiles identical to non-D_2_O groups (Fig.5c, d). These results indicate that lignin degraders incorporate deuterium from D_2_O into biomolecules via metabolic activity, whereas non-degraders lack deuterium assimilation pathways due to their inability to utilize lignin.

**Figure 5.**
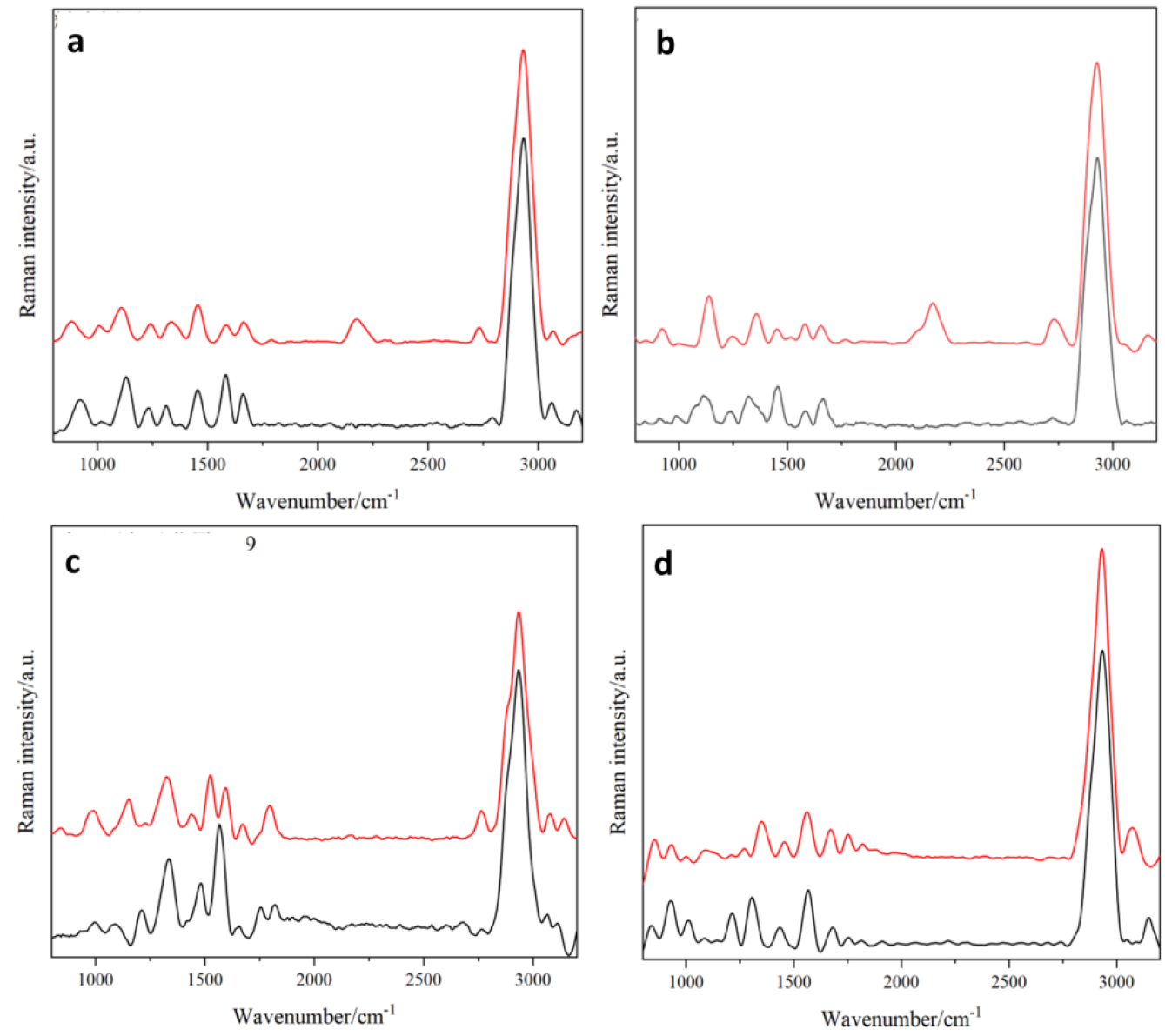
Raman spectra and %C-D ratios of lignin-degrading strains (a) *Pleurotus ostreatus* G5, (b) *P*.*eryngii* PB and non-ligninolytic strains (c) *Kluyvera sp*. pb29, (d) *Enterobacter sp*. NC02, cultured in media supplemented with alkali lignin and 30% D_2_O, with 0% D_2_O as the control group.

Lignin degradation capacity was quantified using %C-D (C-D band intensity relative to C-D + C-H intensities [26]). PB showed the lowest average %C-D, indicating minimal activity. In contrast, QH demonstrated high %C-D values, highlighting its potential for lignin degradation applications.

This D_2_O-labeling Raman approach provides non-destructive, real-time detection without radioactive tracers, establishing an efficient platform for functional screening of environmental microbes. The presence/absence and intensity of C-D peaks validate metabolic activity and offer quantitative screening criteria for strain selection.

## 4 Discussion

Current strategies for functional screening of live single cells directly from microbiomes remain limited. For instance, fluorescence-activated cell sorting (FACS) encapsulates single cells in droplets, isolates pure cultures through cultivation, and screens metabolic activity via fluorescent probes [27]. However, its broad application is hindered by reliance on fluorescent labeling, functional assessment under pure culture rather than in situ conditions, and low post-labeling cell viability.

This study proposes a “screen-before-culture” approach—single-cell Raman-activated sorting coupled with cultivation (scRACS-Culture)—based on single-cell Raman spectroscopy (SCRS). This method enables in situ metabolic screening prior to pure culture isolation without fluorescent probes. Moreover, SCRS assesses a broader spectrum of metabolic activities compared to fluorescent signals, with continuous technological expansions [28]. Thus, this label-free, in situ phenotyping strategy offers greater universality for functional mining in microbial communities.

A critical challenge for Raman-activated cell sorting (RACS) lies in maintaining cell viability during SCRS detection and sorting, as prolonged laser exposure (particularly for non-resonant Raman peaks like C-D bonds) may compromise cellular integrity [29]. To address this, we developed a Raman-activated sorting chip integrating SCRS detection, cell sorting, droplet encapsulation, and single-cell export functions. This system ensures full viability retention even during non-resonant peak-based sorting (e.g., C-D at 2040–2300 cm^−1^). Applying scRACS-Culture, we successfully isolated lignin-degrading strains and validated their degradation efficacy.

In summary, scRACS-Culture provides an in situ functional screening framework to directly assess and mine cultivable/uncultivable functional microbes from environmental samples. This high-throughput approach eliminates time-consuming pre-cultivation steps and avoids false screening inherent to traditional methods. By enabling non-invasive, probe-free detection of diverse metabolic phenotypes, SCRS-based sorting holds broad applicability, significantly advancing functional single-cell technologies in microbiome science and industrial applications.

